# Cultivation and Sequencing Reveal Nutrient-Dependent Bacterial Responses in Public Restrooms

**DOI:** 10.64898/2026.08.23.746590

**Authors:** Jieruiyi Weng, Bei-Wen Ying

**Affiliations:** School of Life and Environmental Sciences, University of Tsukuba, 1-1-1 Tennodai, Tsukuba, 305-8572 Ibaraki, Japan; MiCS, University of Tsukuba, 1-1-1 Tennodai, Tsukuba, 305-8572 Ibaraki, Japan

**Keywords:** bacterial diversity, microbial community, resource availability, culturable bacteria, public sanitation

## Abstract

Microbial communities in indoor environments are shaped by resource availability and disturbances, yet their growth dynamics and compositional changes remain unclear. Here we combined quantitative colony growth analysis with 16S rRNA gene sequencing to investigate bacterial communities on public restroom surfaces before and after routine cleaning under varied nutrient conditions. Cultivation revealed that nutrient availability strongly influenced bacterial growth and selectively enriched distinct taxa, while cleaning caused limited shifts in overall community structure and diversity. Correlations between growth parameters and diversity indices were weak, indicating that taxon-specific responses to nutrients primarily drive growth outcomes. These findings suggest that resource composition, rather than cleaning disturbance, governs bacterial growth and community assembly in built environments. Integrating culture-based phenotyping with sequencing provides a comprehensive framework to understand microbial dynamics following environmental perturbations.

## Introduction

As modern society advances, people are increasingly indoor-bound, with about 90% of their time spent indoors [1, 2]. Due to this prolonged exposure, the indoor environment, especially its bacterial communities, is increasingly impacting human health [3–6]. Indoor bacterial communities have gained more attention because they are abundant and complex [7, 8], with notable differences across various spatial scales [9, 10]. These differences are influenced by geographic location [11, 12], climate [13], and architectural characteristics [14]. Even within the same indoor environment, distinct bacterial communities can vary significantly across different functional spaces [15]. Studies have found that toilets typically harbor the most diverse and complex bacterial communities [15, 16], which makes toilets a particularly important target for microbial research. Intensive studies on toilet bacterial communities have primarily aimed at identifying the types of bacteria present [17–19] and detecting specific pathogens [20, 21]. Thus, the cleanliness of toilets is often studied in association with the bacterial communities.

The impact of cleaning on bacterial communities has become a significant concern [22–24], particularly in public urinals and toilets [25–27]. To ensure sanitary conditions and minimize microbial transmission risks in public restrooms, routine cleaning is commonly practiced [23, 28]. In general, cleaning has been viewed mainly as a way to reduce microbial loads [29, 30]. However, microbial ecology insights indicate that cleaning should be viewed as an environmental disturbance rather than merely a removal process [31, 32]. By altering nutrient availability [33], water flow [34], and the surface microenvironment [35], cleaning has the potential to influence the diversity of the bacterial communities inhabiting surfaces. Certain resistant taxa might survive or become more abundant after cleaning, which could alter the community structure [36, 37]. More studies that directly compare microbial communities before and after cleaning are required to offer quantitative insights into how this disturbance affects community structure, diversity, and functional potential in toilet environments, as such data are still scarce.

To understand environmental microbiomes, 16S rRNA sequencing is commonly used [38, 39]. In our previous study, we applied culture-independent 16S rRNA gene sequencing to investigate bacterial communities inhabiting public urinals before and after routine cleaning and demonstrated that cleaning caused only limited changes in overall bacterial diversity despite detectable shifts in bacterial composition and predicted community functions [40]. However, because sequencing-based analyses provide only compositional information, it remained unclear whether the bacterial populations detected after cleaning retained similar growth capacities or responded differently to changes in nutrient availability. Because it effectively characterizes microbial diversity but cannot differentiate between living and dead cells [41, 42] or growth potential [43]. Interpreting sequencing data directly as indicating only live bacteria may result in misleading conclusions. As a considerable proportion of bacteria may be dormant or dead [44], direct sequencing is insufficient to reflect the truly active bacterial communities in the environment. To address this limitation, culture-based methods can provide complementary information by focusing on viable bacteria [45, 46] and enabling the quantitative assessment of phenotypic characteristics such as colony growth and expansion dynamics [47, 48]. Although culture-dependent approaches are less frequently incorporated into environmental microbiome studies, combining cultivation with 16S rRNA sequencing makes it possible to directly relate bacterial community composition to colony growth responses.

Building upon our previous culture-independent investigation of public restroom bacterial communities [40], the present study focused on the culturable fraction of these environmental communities. By integrating quantitative colony growth analysis with 16S rRNA sequencing of cultured bacteria, we investigated how cleaning-associated disturbance and nutrient availability influence bacterial growth responses and community composition. This integrated approach offers novel insights into the ecological mechanisms that connect environmental bacterial community composition with colony growth dynamics in built environments.

## Materials and Methods

### Sampling, Culturing, and Imaging Dataset

Both the culture-independent environment samples 16S rRNA sequencing dataset and the plate culture growth datasets analyzed in this study were derived from our previously published public toilet disturbance experiment [40]. Detailed protocols regarding surface swabbing, media formulations (LB, 1% LB, M63, MIX, and 10% MIX), plate inoculation (100 μL suspension per plate), 20-day incubation, daily CCD imaging (ATTO AE-6932GXES), and ImageJ-based colony parameter calculations (*C_max_*, *R_max_*, *v_max_*, and *a_max_*) were fully reported in our prior study. While the initial work collected data across 11 urinals (220 agar plates), the present study specifically analyzed a targeted subset of 140 agar plates derived from 7 representative urinals across all 6 sampled public rest areas, summarized in Table S1 as culturable bacterial communities. This subset selection was guided by two criteria: (i) Spatial coverage, ensuring that all six geographically distinct sampling locations were represented to preserve total environmental diversity; and (ii) availability of sufficient bacterial biomass for standardized DNA extraction and high-throughput sequencing. This subset enabled direct integration of colony growth dynamics with corresponding culture-dependent bacterial community profiles obtained by 16S rRNA gene sequencing.

### DNA purification and 16S rRNA sequencing

For cultured samples, all formed colonies on the plate were collected using a sterilized toothpick and transferred into a sterilized 1.5 mL microtube (WATSON). One microtube was used for two plates under the same sample and culture conditions. The microtubes comprising the colonies were stored at -80 for future use. The cell lysates and DNA purification were performed according to our previous studies [49, 50] with slight modifications. 480 μl of 50 mM EDTA (UltraPure™ 0.5M EDTA, pH 8.0, Thermo Fisher Scientific) and 120 μl of 10 mg/ml lysozyme solution (Lysozyme from chicken egg white, Sigma) were added to the microtube comprising the colonies. The colony lysates were incubated at 37□ (BioShaker BR-43FM, TAITEC) for 40 min and centrifuged at 16,000 g (3520, KUBOTA) for 2 min. The supernatant was removed, and the pellet was subjected to DNA purification. 600 μl nuclei lysis solution (Promega) was added and incubated at 80□ (BLOCK INCUBATOR BI-535A, ASTEC) for 5 min. After cooling the solution on ice for 5 min, 600 μl PhOH/CHCl3/Isoamyl alcohol (25:24:1) (Phenol:Chloroform: Isoamyl Alcohol 25:24:1 Saturated with 10 mM Tris, pH 8.0, 1 mM EDTA, Sigma) was added and well mixed. The mixed solution was centrifuged at 16,000 g, 4°C, for 5 min, and the supernatant was collected for DNA purification according to the manufacturer’s instructions (Wizard® Genomic DNA Purification Kit, Promega). 30 μl DNA-Rehydration solution (Wako) was added and incubated at 65□ for one hour to resuspend the purified DNA. The final DNA solution was stored at 4°C. The purified DNA samples were sent out for 16S rRNA sequencing (Bioengineering Lab. Co., Ltd.). The V4 regions of all samples were sequenced using the universal primer pair 515F (5′-GTGCCAGCMGCCGCGGTAA-3′) and 806R (5′-GGACTACHVGGGTWTCTAAT-3′) [51]. The sequencing data were deposited on *figshare* under the accession number 31329844.

### Differential abundance analyses

Raw sequences were processed using the QIIME 2 pipeline (version 2022.8). Sequence denoising, including the removal of chimeric and noisy reads, was performed using the DADA2 plugin to generate the amplicon sequence variant (ASV) table and representative sequences. Taxonomic assignment was conducted using the feature-classifier plugin by aligning representative sequences against the Greengenes database (version 13.8) at 97% identity. Phylogenetic trees were constructed using the alignment and phylogeny plugin. Raw sequences were processed by the Bioengineering Lab. Co., Ltd. To facilitate cross-batch comparisons, the ASV tables and representative sequences from independent runs were integrated using R (v4.5). Sequence manipulation and cross-referencing were performed via the ‘Biostrings’ and ‘dplyr’ packages. All sequences were compared based on 100% identity; those with identical sequences and taxonomic annotations across different batches were collapsed into a single, unified ASV, with their respective abundances summed. The integrated ASV set was then re-indexed (e.g., ASV1, ASV2) to serve as a standardized reference for all downstream analyses. The ASVs are summarized in Table S2 as culturable bacterial communities, respectively. A unified phylogenetic tree was reconstructed using the integrated representative sequences. Multiple sequence alignment (MSA) was performed using the ‘msa’ package (utilizing the MUSCLE algorithm). The resulting alignment was converted to a DNAbin object, and a genetic distance matrix was calculated based on the Kimura 2-parameter (K80) model. The phylogenetic tree was subsequently inferred using the Neighbor-Joining (NJ) method via the ‘ape’ package. This unified tree provided the basis for calculating phylogenetically informed metrics, such as UniFrac distances, to assess community dissimilarity. Annotations of ASVs identified in the culturable bacterial communities are summarized in Table S3.

### Bacterial diversity analyses

α diversity was calculated at the genus level based on the ASV abundance table using R(v4.5). Diversity indicators, including Shannon and Simpson indices, were calculated using the diversity() function in the R package ‘vegan’, and the Chao1 richness estimator was calculated using the estimateR() function from the same package. Pielou’s evenness was calculated manually as:

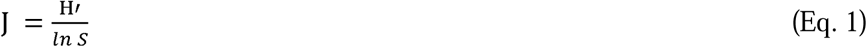

where H′ represents the Shannon diversity index, and S represents the observed ASV richness, which was estimated using the specnumber() function in the R package ‘vegan’ [52]. β diversity analysis was conducted at the genus level based on the amplicon sequence variant (ASV) table, sample metadata, and phylogenetic trees. Data were processed in R (v4.5) using the packages ‘phyloseq’, ‘vegan’, ‘ape’, ‘phangorn’, and ‘pairwise Adonis’. ASV abundances were aggregated to the genus level according to taxonomic annotations, and a genus-level phyloseq object was constructed by integrating abundance data, sample metadata, and a pruned phylogenetic tree. To minimize sequencing depth bias, samples were rarefied to the minimum library size using the rarefy_even_depth() function in the ‘phyloseq’ package. Phylogenetic beta diversity was assessed using weighted UniFrac distances calculated with the UniFrac() function in ‘phyloseq’. Principal coordinate analysis (PCoA) was performed using the ordinate() function to visualize community dissimilarities among samples, and group dispersion patterns were illustrated with 95% confidence ellipses using ggplot2. Permutational multivariate analysis of variance (PERMANOVA) was conducted using the adonis2() function in the ‘vegan’ package to evaluate the effects of environmental factors (e.g., urinals, sampling sites, media, and cleaning conditions) on microbial community composition. Homogeneity of group dispersions was assessed using betadisper() followed by permutation tests (permutest()). For paired before-and-after-cleaning comparisons, PERMANOVA was performed with sampling location specified as a stratification factor. Additional pairwise PERMANOVA comparisons among groups were conducted using repeated adonis2() tests with false discovery rate (FDR) correction applied to multiple comparisons. False discovery rate (FDR) correction was also applied, where appropriate, to other multiple-comparison analyses, including paired Wilcoxon tests and differential abundance analyses presented in volcano plots.

### Enriched bacterial taxa analyses

Differential abundance analysis was performed at the genus level using DESeq2 implemented in R (v4.5). The ASV abundance table was first filtered to remove chloroplast and mitochondrial sequences, and taxa without genus-level annotation were excluded. ASV counts were aggregated to the genus level using taxonomic agglomeration. Only cultured samples from LB, M63, and MIX conditions were retained for subsequent analysis. Pairwise comparisons were performed among the three culture conditions (LB vs M63, LB vs MIX, and M63 vs MIX). For each comparison, only matched samples collected from the same urinal were included to maintain the paired experimental design. Genera with fewer than 10 reads in at least two samples were removed prior to differential abundance analysis. DESeq2 models were fitted using a negative binomial generalized linear model with the design formula ∼ Urinal + Condition, where urinal identity was included as a blocking factor to account for repeated measurements. Size factors were estimated using the positive-counts method, and dispersion parameters were estimated using the local fitting approach. Differentially abundant genera were identified based on log2 fold changes and Benjamini–Hochberg adjusted p-values (FDR < 0.05). Significant genera detected across pairwise comparisons were further summarized according to their recurrence among comparisons.

### Variation contribution analyses

A contribution score for each ASV to evaluate the relative importance of variation was calculated accordingly (Eq. 2), based on our previously established mathematical framework.

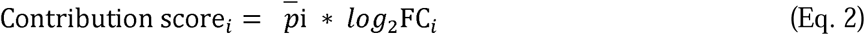

where *p̅*_i_ represents the mean relative abundance of the i_th_ ASV across all samples (calculated after per-sample normalization), and log_2_*FC_i_*denotes its differential abundance (from DESeq2 analysis). Only ASVs with significant differential abundance (adjusted *p* < 0.05) were retained for contribution score calculation. ASVs were then ranked according to their contribution scores to identify the major drivers of structural differences between groups. All analyses were performed in R (v4.5) using the packages dplyr, readr, tidyr, ggplot2, and forcats.

## Results

### Nutrient availability shapes bacterial community growth

To investigate how physical perturbation and nutrient availability influence bacterial growth dynamics, we first quantified the overall community-level growth responses across five culture media before and after cleaning. No significant differences in fold change before and after cleaning were detected for any of the growth parameters (Fig. 1A), indicating that the response of bacterial community growth to cleaning remained consistent across different nutrient conditions. To characterize the effects of nutrient availability on colony growth, the absolute values of the four growth parameters were compared among the five culture media before and after cleaning (Fig. S1). Significant differences were observed among media for all growth parameters under both conditions (Kruskal–Wallis test, all P < 0.01). LB consistently supported the greatest colony abundance and expansion, followed by M63, whereas 1% LB, MIX, and 10% MIX exhibited substantially lower growth. Importantly, these medium-dependent growth patterns were preserved after cleaning. Thus, although nutrient availability strongly influenced colony growth, cleaning did not modify these medium-dependent differences. In other words, although bacterial growth differed among media, the magnitude of the response to environmental perturbation was largely independent of nutrient availability.

**Figure 1.**
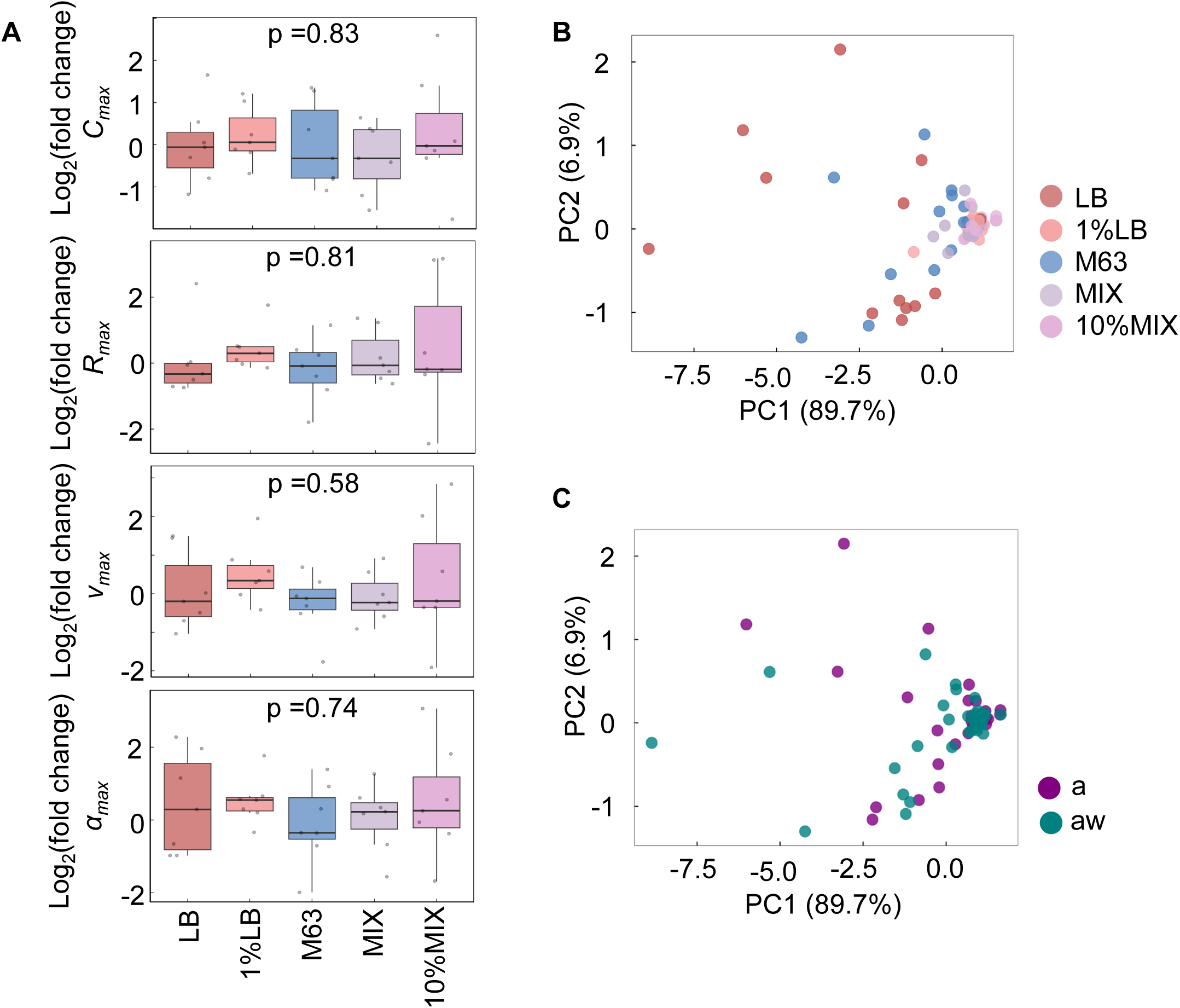
Growth analyses of culturable bacteria on different media. **A.** Comparison of fold changes in colony growth parameters among different media. Boxplots show the fold changes in *C_max_*, *Rmax*, *vmax,* and *amax* for the five media (LB, 1%LB, M63, MIX, and 10%MIX). Each point represents one biological replicate (n = 7 urinals), with the mean value of two technical replicates obtained from the same urinal under the same culture condition. Boxes indicate the interquartile range (IQR), the center line represents the median, and whiskers extend to 1.5 × IQR. The overall effect of culture medium was evaluated using a linear mixed-effects model, with culture medium as a fixed effect and urinal identity as a random effect. Principal component analysis (PCA) of colony growth parameters before and after cleaning. PCA was performed separately using the four colony growth parameters measured under the five culture media before **(B)** and after **(C)** cleaning. Prior to PCA, all variables were standardized to zero mean and unit variance. Each point represents one biological sample (7 urinals × 5 media), and the percentages shown on the axes indicate the proportion of variance explained by each principal component.

To further evaluate whether bacterial communities could be differentiated based on their overall colony growth characteristics, principal component analysis (PCA) was performed using the four growth parameters (*C_max_*, *R_max_*, *v_max_*, and *a_max_*). The first principal component (PC1) explained 89.7% of the total variance, whereas the second principal component (PC2) explained 6.9%. Before cleaning, samples cultured on LB and M63 tended to separate from those cultured on the other media along PC1; however, substantial overlap remained among media, and no discrete clusters were observed (Fig. 1B). Likewise, samples collected before and after cleaning largely overlapped in PCA space (Fig. 1C), indicating that routine cleaning did not induce a consistent shift in the overall colony growth characteristics of the bacterial communities. Together, these findings suggest that bacterial community growth was robust to cleaning, whereas nutrient availability primarily influenced the magnitude of colony growth while preserving the overall multivariate growth characteristics of the bacterial communities.

### Cultivation alters bacterial community composition but not overall diversity

To determine how cultivation under different nutrient conditions influenced bacterial community composition, 16S rRNA amplicon sequencing was performed on the cultured communities before and after cleaning. A total of 40 plates comprising three media (LB, M63, and MIX) were subjected to sequencing analysis (Fig. 2A). A total of 208 ASVs were identified from 40 culture plates representing three media (LB, M63, and MIX) and mainly annotated to five bacterial phyla, with *Proteobacteria*, *Actinobacteria*, and *Firmicutes* being the dominant phyla, accounting for approximately 63.1%, 19.0%, and 14.9% of the total sequencing reads, respectively (Fig. 2B, Table S4). The changes in the bacterial compositions before and after cleaning were further evaluated at the family level (Table S5). A large proportion of bacterial families was shared between samples collected before and after cleaning in the cultured (Fig. 2C) communities, indicating that cleaning caused only limited changes in overall community composition. To further investigate how cultivation under different nutrient conditions influenced bacterial community composition, the taxonomic profiles of the cultured communities were compared with those of the corresponding culture-independent environmental communities reported in our previous study [40]. In contrast, cultivation substantially reduced taxonomic richness, with the total number of detected families decreasing from 170 in the original communities to 57 after cultivation, reflecting selective enrichment during cultivation. Interestingly, four bacterial families were detected exclusively in the cultured communities (Fig. 2D), suggesting that cultivation enabled the recovery of a small number of low-abundance or otherwise undetected taxa from the original communities.

**Figure 2.**
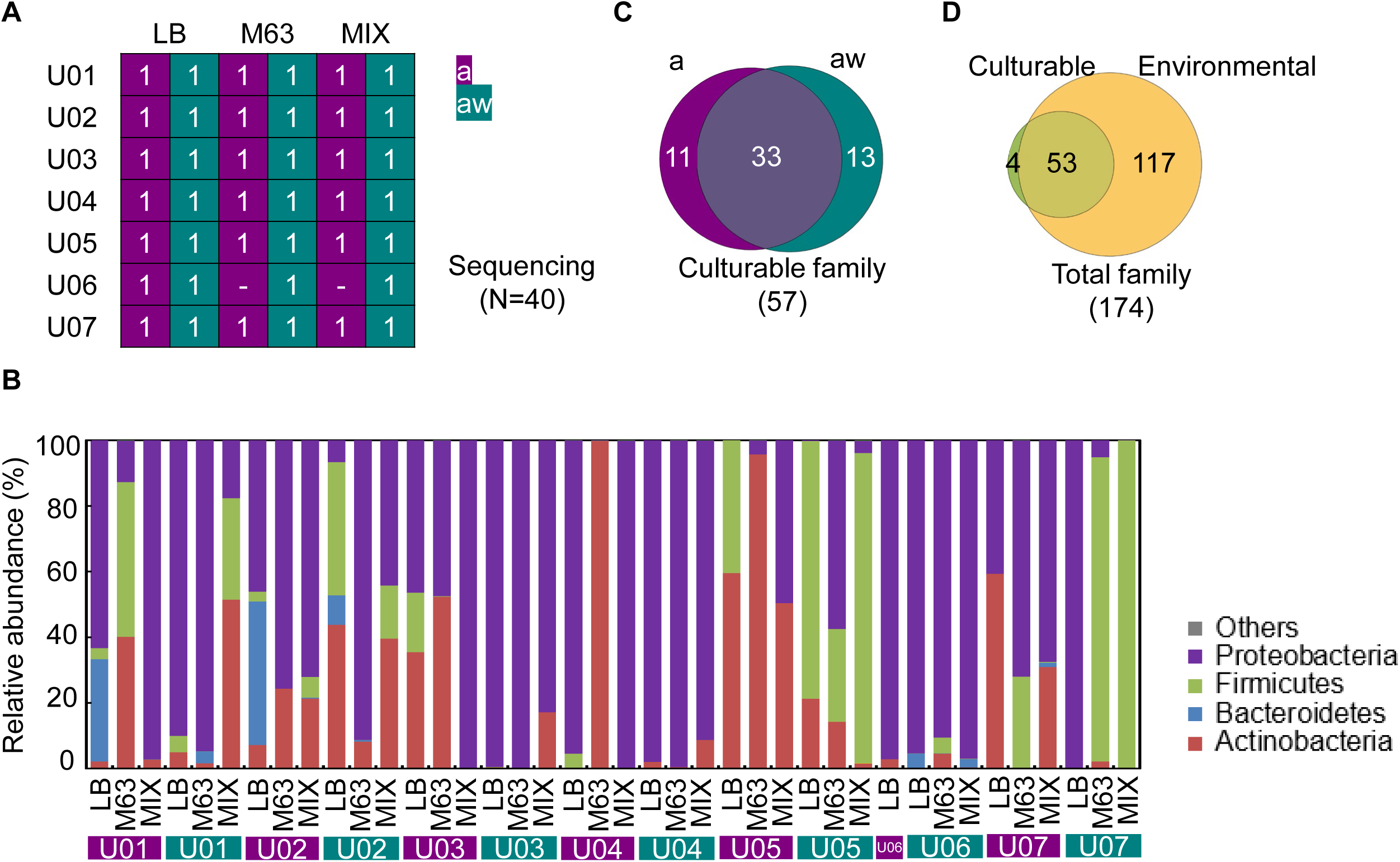
Bacterial communities and culturable bacterial communities inhabiting urinals. **A.** Plates used for sequencing. The urinal ID, medium, cleaning condition, and count are indicated. **B.** Compositions of culturable bacterial communities. Color variation indicates the different phyla. The urinal ID, medium, and cleaning condition are indicated. **C.** Venn diagram of culturable bacterial taxa before and after cleaning. **D.** Venn diagram of culturable and environmental bacterial taxa. The number of taxa at the family level is indicated.

To further assess whether cleaning or nutrient conditions affected the diversity and overall structure of bacterial communities after cultivation, we performed α- and β-diversity analyses at the genus level, incorporating culture-independent environmental samples from our previous study [40]. Before cleaning, environmental samples showed significantly higher Shannon, Simpson, and Chao1 indices than bacterial communities cultured on LB, M63, and MIX media, while Pielou’s evenness was similar between environmental samples and most cultured communities (Fig. 3A). This pattern persisted after cleaning (Fig. 3B), indicating that routine cleaning did not alter the relationship between cultured and original communities. Direct comparisons before and after cleaning revealed no significant differences in α-diversity for any culture condition, including LB, M63, MIX, and environmental samples (Fig. 3C). Among cultured communities, α-diversity was comparable across the three media both before and after cleaning (Fig. S2), suggesting that nutrient composition did not substantially affect overall α-diversity. Consistent with these findings, Bray--Curtis PCoA showed that significant differences in community composition were primarily due to the separation between cultured and environmental samples, both before (Fig. 3D) and after cleaning (Fig. 3E), while bacterial communities recovered on LB, M63, and MIX largely overlapped. When analysis was limited to cultured communities, no significant differences in β-diversity were detected among the three media, either before or after cleaning (Fig. S3).

**Figure 3.**
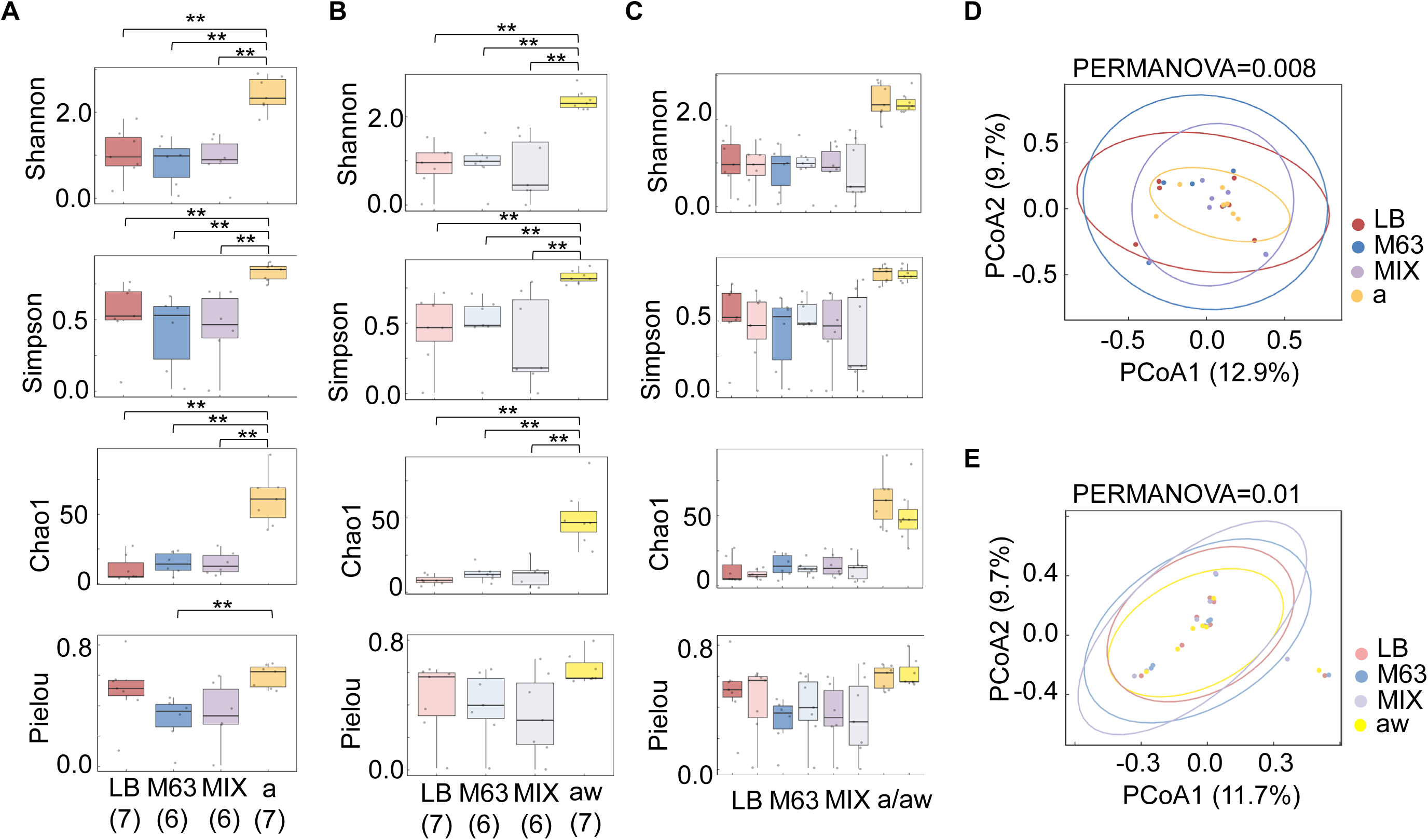
Genus level bacterial diversity on different media. **A.** Bacterial α diversity on the three media and environment samples before cleaning. **B.** Bacterial α diversity on the three media and environment samples after cleaning. **C.** Comparison of bacterial α-diversity before and after cleaning for each culture condition. Principal coordinate analysis (PCoA) based on genus-level Bray–Curtis dissimilarities of three media and environment samples before (D) and after (E) cleaning. Samples were rarefied to an equal sequencing depth prior to analysis. Each point represents one sample, and ellipses represent the 95% confidence ellipses for each culture condition. The percentages shown on the axes indicate the proportion of total variation explained by each principal coordinate. Differences in community composition among culture conditions were assessed using PERMANOVA based on Bray–Curtis dissimilarities.

Together, these results indicate that cultivation consistently distinguished cultured communities from the original environmental microbiota, whereas routine cleaning and differences in nutrient composition among the tested culture media had only limited effects on the overall diversity and community structure of the cultured bacterial communities. These observations suggest that the pronounced differences in colony growth among culture media (Fig. S1) were not accompanied by corresponding changes in overall bacterial diversity or community composition.

### Relationships between bacterial diversity and colony growth parameters

To further investigate whether variation in bacterial α-diversity was associated with colony growth characteristics, Spearman correlation analyses were performed between genus-level α-diversity indices and colony growth parameters (*C_max_*, *R_max_*, *v_max_*, and *a_max_*) before and after cleaning. Because colony growth responses differed among culture media, the analyses were performed separately for each medium to determine whether associations between bacterial diversity and colony growth were influenced by nutrient conditions. Before cleaning, only a few significant associations were detected. In LB medium, Chao1 richness showed a significant positive correlation with *R_max_*, whereas no significant correlations were observed in M63 medium, and under MIX conditions, Pielou’s evenness was negatively correlated with both *v_max_* and *a_max_* (Fig. 4A), indicating that communities with higher evenness tended to exhibit reduced colony growth dynamics under this nutrient condition. After cleaning, the correlation pattern again differed among media. The greatest number of significant associations was observed in the LB medium. Specifically, Chao1 richness was positively correlated with *C_max_* and also showed positive correlations with *v_max_* and *a_max_*. Shannon diversity was likewise positively associated with *v_max_* and *a_max_*. In contrast, no significant correlations were detected in M63 or MIX media, although several weak positive and negative trends remained (Fig. 4B). These observations indicate that associations between bacterial diversity and colony growth were context-dependent, varying with both nutrient conditions and cleaning status rather than being consistently conserved across culture media.

**Figure 4.**
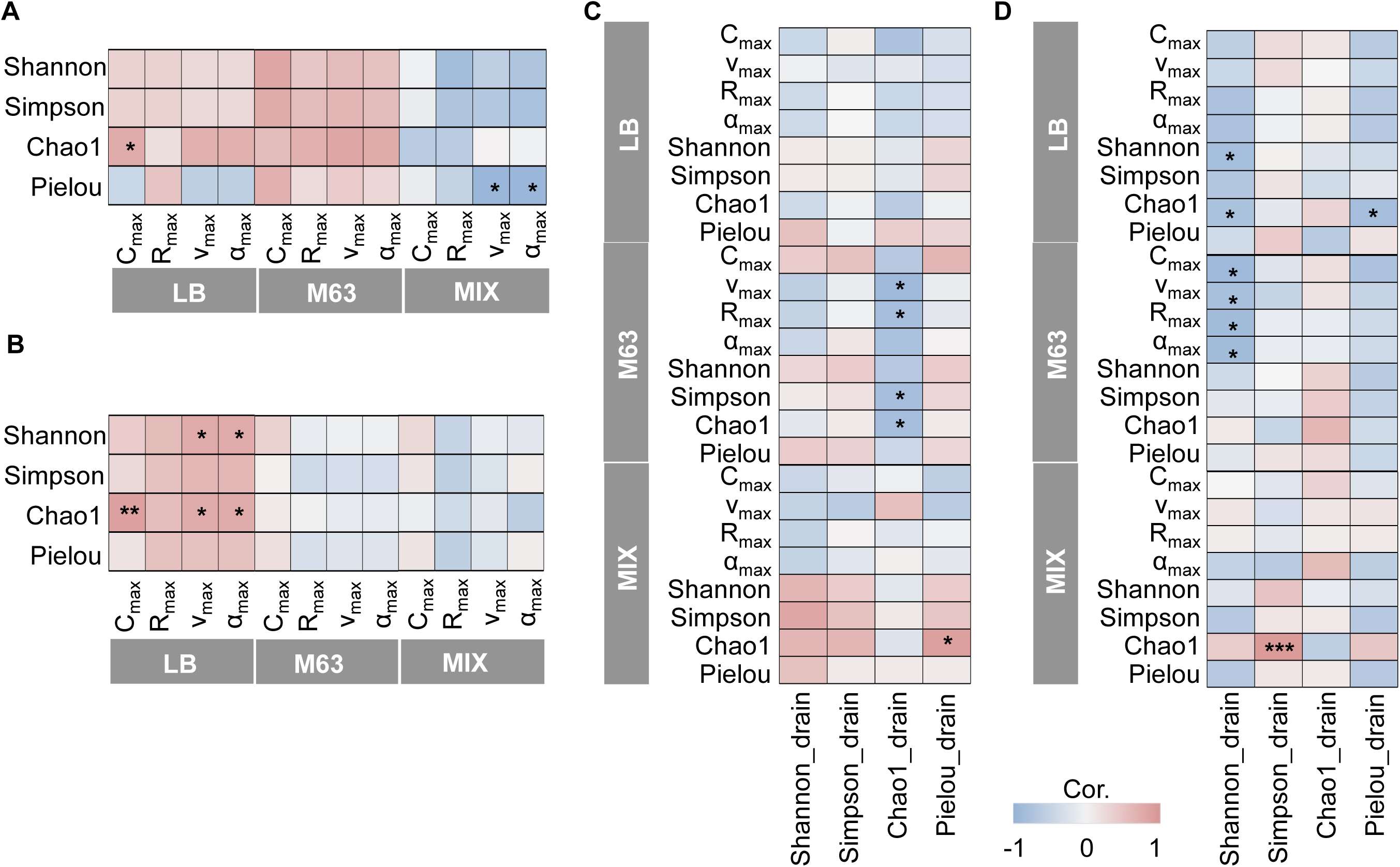
Correlations between colony growth parameters and bacterial diversity. Spearman correlation heatmaps showing the relationships between colony growth parameters (*C_max_*, *Rmax*, *vmax,* and *amax*) and genus-level α diversity (Shannon, Simpson, Chao1, and Pielou) indices under LB, M63, and MIX media conditions before (A) and after (B) cleaning. Heatmaps of the relationships between environment samples’ genus-level α diversity indices and the genus-level diversity of cultured bacteria (Shannon, Simpson, Chao1, and Pielou) as well as colony growth parameters (*C_max_*, *Rmax*, *vmax,* and *amax*) under LB, M63, and MIX media conditions before (C) and after (D) cleaning. Color intensity indicates the Spearman correlation coefficient (ρ), with red and blue representing positive and negative correlations, respectively. Asterisks denote statistically significant correlations (*P < 0.05, **P < 0.01, ***P < 0.001).

To further examine whether the initial diversity of the original environmental bacterial communities was associated with the characteristics of the resulting cultured communities, we analyzed correlations between the diversity metrics of the culture-independent environmental communities reported previously [40] and those of the corresponding cultured communities. Before cleaning, correlations between the α-diversity of the original environmental bacterial communities and the diversity or growth characteristics of cultured bacteria were generally weak (Fig. 4C). Only a few statistically significant correlations, primarily in the Chao1 metric, were detected under individual cultivation conditions, primarily in the M63 medium, while no consistent relationship was observed across the three media. After cleaning, statistically significant correlations were primarily in Shannon (Fig. 4D). However, these significant associations remained sporadic and varied among the cultivation media, with no common pattern across LB, M63, and MIX. Overall, the α-diversity of the original environmental bacterial communities was not consistently associated with either the diversity of cultured communities or colony growth characteristics. Taken together, these results indicate that associations between bacterial diversity and colony growth were context-dependent rather than universal. Although significant relationships were detected under specific nutrient conditions, these associations were not consistently maintained across culture media or before and after cleaning, suggesting that variation in colony growth characteristics cannot be reliably predicted from overall bacterial α-diversity alone, despite the marked differences in colony growth observed among culture media.

### Differentially abundant bacterial taxa among culture media

Although overall bacterial α-diversity showed only limited and context-dependent associations with colony growth (Fig. 4), colony growth differed markedly among culture media (Fig. S1). Therefore, differential abundance analysis was performed to determine whether specific bacterial taxa exhibited medium-dependent shifts in abundance despite relatively stable community-level diversity and composition. Before cleaning, relatively few taxa differed significantly among the three culture media (Fig. 5A), indicating limited medium-dependent taxonomic selection under baseline conditions. In the comparison between LB and M63, *Micrococcus* was enriched in LB, whereas *Kocuria* was enriched in M63, indicating that these genera responded differently to nutrient conditions. Comparisons involving MIX medium further identified enrichment of *Agrobacterium* relative to the alternative media. Overall, only a small number of taxa exhibited significant differential abundance before cleaning. In contrast, substantially stronger medium-dependent selection was observed after cleaning (Fig. 5B). Compared with M63, LB preferentially enriched *Blastomonas* and *Sphingomonas*, whereas M63 showed significant enrichment of *Methylobacterium* and an unclassified member of the family *Phyllobacteriaceae*. Comparisons between M63 and MIX further identified *Acinetobacter*, *Burkholderia*, and *Phyllobacteriaceae* as significantly enriched in M63. *Blastomonas* showed the largest estimated fold change among the detected taxa (absolute log2 fold change = 18.8), although this difference was driven by strong condition-specific occurrence patterns. Collectively, these results demonstrate that nutrient availability selectively enriched distinct bacterial taxa despite the relatively limited differences observed in overall bacterial diversity and community composition. Moreover, a greater number of differentially abundant taxa were detected after cleaning than before cleaning, indicating that medium-dependent taxonomic differentiation became more pronounced following cleaning.

**Figure 5.**
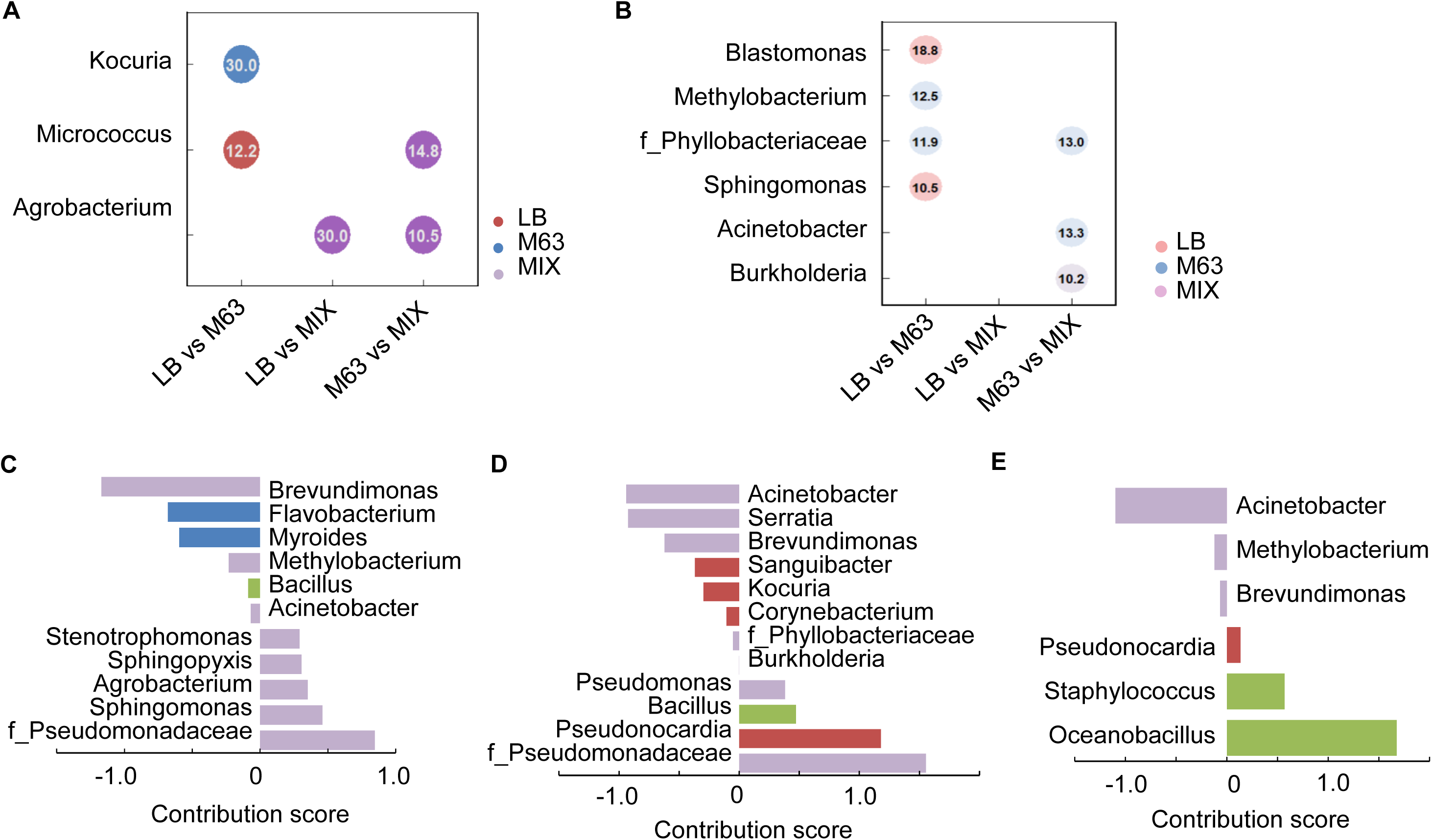
Differentially enriched bacterial taxa among culture media before and after cleaning. Bubble plots showing bacterial taxa that were significantly differentially abundant among the three culture media (LB, M63, and MIX) before (A) and after (B) cleaning, as identified by DESeq2 analysis. Pairwise comparisons were performed for LB vs M63, LB vs MIX, and M63 vs MIX using a paired design with urinal identity included as a blocking factor (∼ Urinal + Base). Only taxa with an adjusted P value (Benjamini–Hochberg corrected) < 0.05 are shown. Bubble colors indicate the culture medium in which each taxon was enriched, as determined by the direction of the log2 fold change, and the values displayed inside the bubbles represent the absolute log2 fold change value. Taxa lacking genus-level annotation are labeled using the highest confidently assigned taxonomic rank (e.g., family or order). **C.** Contribution of genera to the cleaning-mediated changes in LB media. **D.** Contribution of genera to the cleaning-mediated changes in M63 media. **E.** Contribution of genera to the cleaning-mediated changes in MIX media. The genera of significant positive or negative contribution scores are shown.

To identify the bacterial taxa contributing most to the cleaning-associated compositional differences within each cultivation medium, contribution scores were calculated by integrating the relative abundance and differential abundance of significantly enriched ASVs. The contribution profiles differed markedly among the three cultivation media, indicating that distinct bacterial taxa accounted for the observed compositional changes under different nutrient conditions. In the LB medium, genera showing positive contribution scores (more abundant after cleaning) were predominantly affiliated with Proteobacteria, whereas the major negative contributors (more abundant before cleaning) included members of Proteobacteria together with a few genera from Bacteroidetes and Firmicutes (Fig. 5C). In the M63 medium, positive contribution scores were distributed across Proteobacteria, Actinobacteria, and Firmicutes, whereas negative contribution scores were dominated by Proteobacteria and Actinobacteria. (Fig. 5D). In the MIX medium, only a limited number of genera contributed to the cleaning-associated differences. Positive contribution scores were mainly associated with Firmicutes, whereas the negative contribution scores were almost exclusively represented by Proteobacteria (Fig. 5E). Overall, different cultivation media exhibited distinct taxonomic contribution patterns, reflecting differences in the bacterial genera associated with cleaning under different nutrient conditions.

Subsequently, the core bacteria were predicted according to previous studies [53–55], at a 75% occurrence frequency (prevalence) and a 1% detection threshold (Fig. S4). This core microbiome analysis was applied comprehensively across both our present cultured datasets and the previously published environmental datasets [40]. Under these stringent criteria, no core genera were identified in the environmental communities before cleaning or in any of the cultured communities, regardless of cleaning status. In contrast, two core genera were detected in the environmental communities after cleaning. These findings indicate that few bacterial taxa were consistently shared among samples, reflecting considerable inter-sample heterogeneity under both environmental and cultured conditions. Overall, these analyses indicate that, although cleaning and nutrient conditions produced only limited changes in overall bacterial diversity (Fig. 3), they selectively altered the abundance of specific bacterial taxa. These findings suggest that differences in colony growth across nutrient conditions were more closely associated with shifts in the abundance of specific bacterial taxa than with broad changes in community diversity. Together with the weak diversity–growth associations observed in Fig. 4, these results indicate that taxon-specific responses to nutrient availability may contribute more strongly to colony growth than overall diversity patterns.

## Discussion

The present study combined community-level cultivation with 16S rRNA gene sequencing to investigate how nutrient availability and environmental disturbance shape bacterial growth in indoor microbial communities. Rather than focusing solely on changes in bacterial community diversity, this study directly linked colony growth dynamics with taxonomic responses under different nutrient conditions, providing ecological insight into the mechanisms underlying bacterial growth in built environments. Compared with the relatively limited effects of cleaning, nutrient availability exerted a much stronger influence on bacterial growth under experimental conditions. These findings suggest that bacterial growth responses were more strongly associated with resource availability than with the environmental disturbance imposed by cleaning (Fig. 1), indicating that bacterial growth responses were more influenced by resource availability than by cleaning. Consistently, comparisons of absolute growth characteristics among media showed substantial variation under different nutritional conditions (Fig. S1), highlighting the importance of resource composition in shaping microbial growth performance. The different media used in this study represented distinct nutritional environments, including nutrient-rich LB, nutrient-limited M63, and the defined MIX medium, together with their diluted formulations. Such differences in nutrient composition can influence bacterial growth by affecting resource availability, metabolic requirements, and physiological strategies of individual bacterial populations [56]. Previous studies have shown that environmental microorganisms often exhibit heterogeneous growth responses depending on nutrient conditions, reflecting differences in resource utilization capacity and metabolic potential among taxa [57, 58]. Therefore, the observed variation in colony growth among cultivation conditions likely reflects differential responses of bacterial populations to available resources rather than a simple consequence of overall community growth capacity.

Cultivation altered the observed bacterial community composition compared with environmental samples, while several dominant bacterial families remained consistently represented across cultivation conditions (Fig. 2). This pattern suggests that cultivation reshaped the relative composition of bacterial communities while retaining some dominant bacterial families from the original environmental microbiota, rather than simply reducing the observable community. Despite the compositional changes introduced by cultivation, a large proportion of bacterial families were shared between samples collected before and after cleaning, consistent with our previous finding that routine cleaning had only limited effects on overall bacterial community composition [40]. This observation suggests that the mild environmental disturbance imposed by cleaning did not fundamentally restructure the bacterial community. Taxa detected exclusively in cultured samples may have been present at low, initially undetectable abundances but became enriched during cultivation; conversely, taxa detected only in environmental samples might have been metabolically inactive, slow-growing, or lacking the physiological capacity to respond to the nutritional environment provided in this study [44].

Our previous study demonstrated that bacterial α-diversity in after cleaning samples was significantly associated with cleaning duration [40], whereas the present study compared bacterial communities collected immediately before and after routine cleaning and detected no significant differences in α-diversity (Fig. 3, Fig. S2). Taken together, these findings suggest that cleaning may induce subtle changes in bacterial diversity; however, these changes are insufficient to produce statistically detectable differences when bacterial communities are compared simply before and after cleaning.

Although several statistically significant correlations were identified, the relationships between bacterial diversity and colony growth varied across nutrient conditions and cleaning states, indicating that these associations were nutrient-dependent rather than universally conserved (Fig. 4). Nevertheless, most growth parameters showed no consistent relationship with bacterial α-diversity, suggesting that bacterial diversity alone was a poor predictor of colony growth. Similar inconsistencies between microbial diversity and bacterial growth responses have been reported previously, where community-level diversity often fails to predict growth performance because bacterial proliferation is more strongly driven by the physiological characteristics and nutrient responsiveness of dominant taxa than by overall community diversity [59, 60]. Furthermore, the diversity of the original urinal bacterial communities was not a reliable predictor of either colony growth or the diversity of cultured communities (Fig. 4), indicating that cultivation outcomes depended more on bacterial responses to the nutrient environment than on the overall diversity of the original environmental microbiota. This inconsistency likely reflects both the selective effects of nutrient availability during cultivation and the fundamental differences between sequencing- and cultivation-based approaches. Whereas 16S rRNA sequencing captures DNA from the entire bacterial community, including inactive or non-growing populations, colony growth reflects the metabolically active bacterial fraction capable of proliferating under the experimental conditions [61, 62]. Therefore, in this study, community diversity alone was not a strong predictor of the growth potential of environmental microbiomes, particularly when microbial communities encountered altered resource conditions that may favor specific physiological strategies.

Differential abundance analysis provided insights into the potential mechanisms underlying the pronounced differences in colony growth observed among cultivation media by revealing that specific bacterial taxa were selectively enriched under different nutrient conditions (Fig. 5). These taxon-specific enrichments may be associated with the variation in colony growth observed among cultivation conditions [63, 64]. This finding is consistent with earlier frameworks showing that nutrient additions drive community assembly via the selective proliferation of responsive subpopulations [65]. Interestingly, a greater number of differentially enriched taxa were detected after cleaning than before cleaning, indicating that bacterial responses to nutrient availability became more differentiated following environmental disturbance. This pattern suggests that environmental disturbance altered the taxonomic composition of the source communities, thereby changing the pool of bacteria available for selection under different nutrient environments. Contribution score analysis further demonstrated that the bacterial genera contributing most to the cleaning-associated compositional differences varied among the three cultivation media. Because contribution scores incorporated both relative abundance and differential abundance, the analysis highlighted bacterial taxa that contributed most substantially to community differences rather than simply those showing statistically significant changes. The distinct contribution patterns observed across cultivation media further support the view that nutrient availability influences bacterial communities through taxon-specific responses rather than uniform shifts across the entire community. In addition, core microbiome analysis revealed that few bacterial taxa were consistently detected across samples under the stringent prevalence threshold (Fig. S4), suggesting that the stability of these bacterial communities was not dependent on a fixed set of core taxa but involved changes in the relative abundance of different bacterial members across environmental conditions.

Taken together, the present findings indicate that differences in bacterial growth among nutrient conditions cannot be explained by broad changes in bacterial community diversity alone. Instead, bacterial growth in built environments appears to be governed primarily by taxon-specific responses to resource availability, whereby different nutrient conditions promote the proliferation of bacterial populations with distinct growth capacities. In this context, cleaning acted as an environmental disturbance that produced only subtle changes in overall community diversity and composition. In contrast, nutrient availability exerted a stronger influence on bacterial growth by modifying the relative abundance of responsive bacterial taxa. These findings suggest that understanding microbial dynamics in built environments requires linking community composition with microbial growth phenotypes rather than relying solely on diversity-based community descriptions. Integrating cultivation-based growth phenotyping with sequencing-based community profiling therefore provides a complementary ecological framework for elucidating how environmental bacterial communities respond to changes in resource availability following disturbance.

Several limitations of the present study should be acknowledged. First, this study was conducted as a pilot investigation based on bacterial communities collected from a limited number of urinals within a restricted geographical region. Although the observed patterns were consistent across the analyzed samples, larger-scale investigations involving more sampling sites and broader geographical coverage will be necessary to determine the generality of the ecological patterns reported here. Second, only a limited number of cultivation media representing different nutrient conditions were examined. Although these media encompassed both nutrient-rich and nutrient-poor environments, they cannot fully represent the diversity of nutritional conditions encountered in built environments. Expanding the range of cultivation media or incorporating additional environmental factors would provide a more comprehensive understanding of how resource availability shapes bacterial growth and community assembly. In addition, cultivation-based analyses inherently capture only the bacterial populations capable of growing under the selected laboratory conditions. Consequently, the observed taxon-specific responses represent the cultivable fraction of the environmental microbiota and may not fully reflect the responses of uncultured bacterial populations. Integrating cultivation with complementary cultivation-independent approaches or expanded culturomics strategies would further improve our understanding of microbial responses to environmental disturbance and resource availability. Finally, the present study identified taxon-specific enrichment as a likely mechanism underlying differences in colony growth; however, the physiological mechanisms responsible for these differential growth responses remain unclear. Future studies combining cultivation experiments with genomic, transcriptomic, or metabolomic analyses may help clarify how individual bacterial taxa respond to changing nutrient environments and environmental disturbances.

## Data Availability

All data generated and analyzed are provided in the supplementary tables and deposited in the public repository, *figshare*, which can be accessed at the following URLs: https://doi.org/10.6084/m9.figshare.31239340 https://doi.org/10.6084/m9.figshare.31329844

## Author contributions

JW performed the experiments, analyzed the data, and drafted the manuscript. BWY conceived the study, validated the data, and wrote the manuscript. All authors approved the final manuscript.

## Acknowledgments

We thank Takamasa Hashizume for analytical support.

