## Supplemental figures and table captions for "Cultivation and Sequencing Reveal Nutrient-Dependent Bacterial Responses in Public Restrooms"

**Supplementary figures (Figure S1~S4)** **p. 2~5**
**Captions for supplementary tables (Table S1~S5)** **p.6**

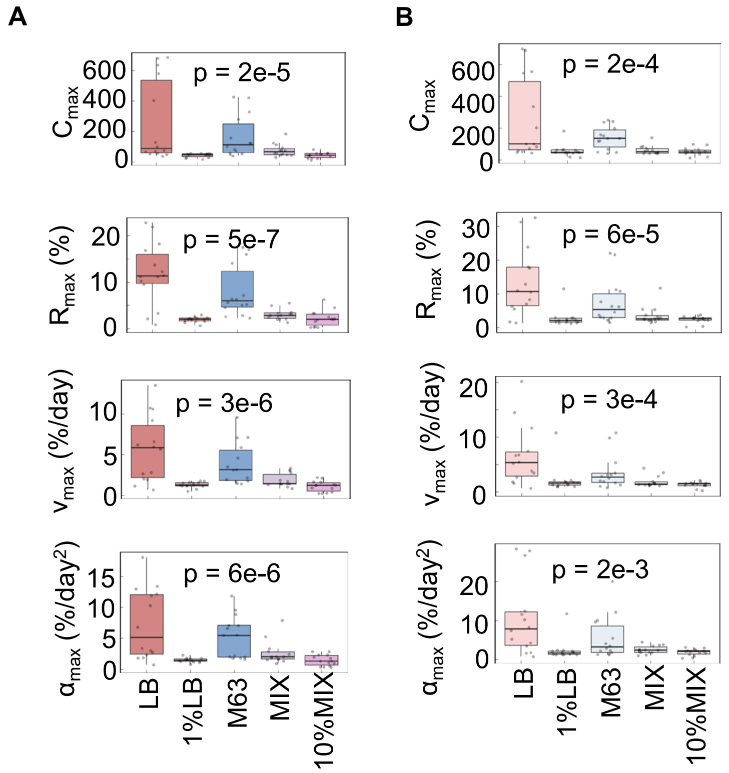


**Figure S1 Growth of culturable bacteria on different media. A.** Growth on five different media before cleaning. **B.** Growth on five different media after cleaning. Four parameters, $C_{max}$, $R_{max}$, $v_{max}$, and $a_{max}$, are shown from top to bottom panels. The p-values of Kruskal–Wallis tests are indicated.

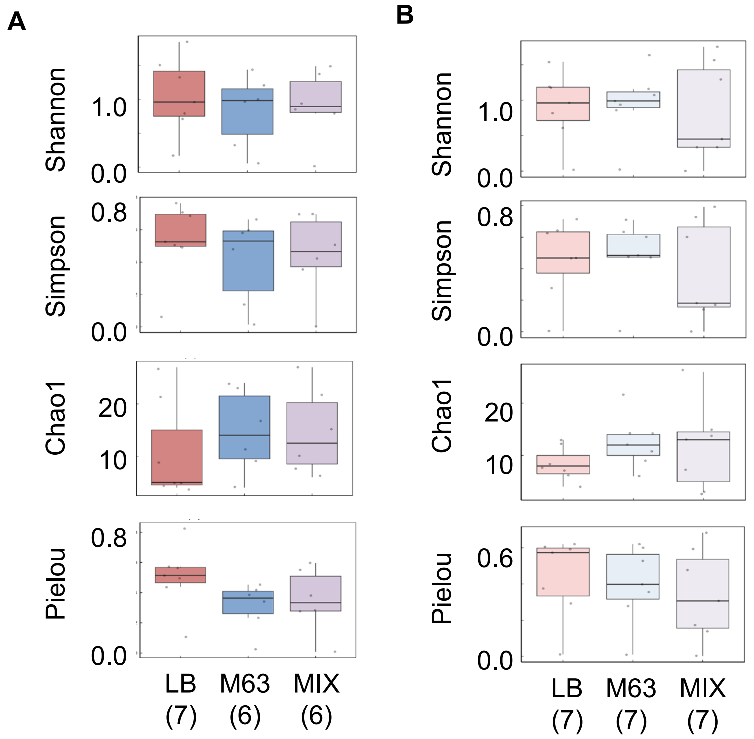


**Figure S2 Bacterial α diversity on the three media before cleaning.** Genus-level bacterial diversity on the three media before (**A**) and after (**B**) cleaning.

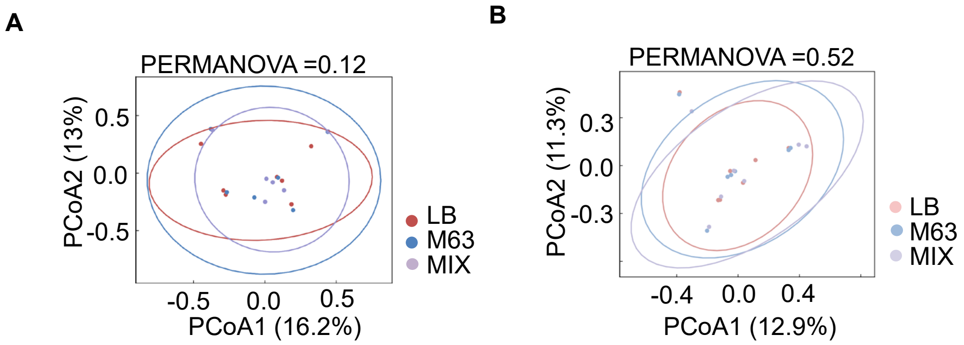


**Figure S3 Bacterial β diversity on the three media after cleaning.** Principal coordinate analysis (PCoA) based on genus-level Bray–Curtis dissimilarities of three media before (**A**) and after (**B**) cleaning.

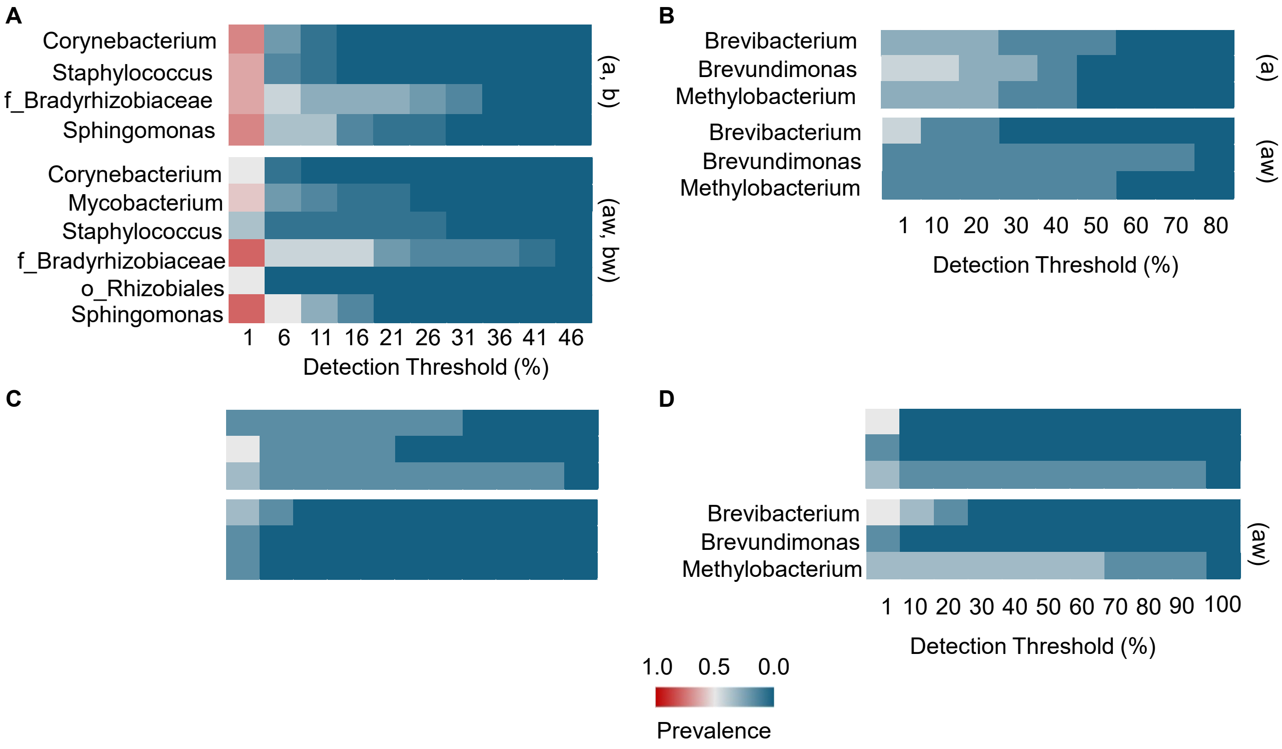


**Figure S4 Core microbiome analysis.** Core microbiomes identified in **A.** environment samples, **B.** LB, **C.** M63 cultures, and **D.** MIX. The upper panels show samples collected before cleaning, whereas the lower panels show samples collected after cleaning. The core microbiome was defined using a prevalence threshold of 75% and a minimum relative abundance (detection) threshold of 1%.

**Supplementary table captions**

**Table S1 Parameters used to evaluate the growth of culturable bacterial communities.** Calculated parameters used for the growth analysis are summarized for each growth medium, cleaning condition, and the IDs assigned to the agar plate, 16S rRNA sequencing, and urinal.

**Table S2 Frequency of ASVs detected in 40 cultured samples.** ASVs used for annotation and diversity analysis are summarized by cleaning condition, growth medium, and the IDs assigned to 16S rRNA sequencing and the urinal.

**Table S3 Annotation of ASVs detected in 40 cultured samples.** Bacterial annotation, from kingdom to species, is provided for each ASV.

**Table S4 Compositions of bacterial communities in 40 cultured samples.** The abundance of each genus within the bacterial community is provided, corresponding to the medium, cleaning condition, and the ID assigned to the urinal.

**Table S5 Changes in bacterial communities caused by cleaning and culture.** Bacterial families detected before and after cleaning in cultured samples are listed, corresponding to Fig. 2C, respectively. The comparison between all bacterial families detected in cultured and environmental samples is also provided, corresponding to Fig. 2D. All labels correspond to the referred figure.
